# Chromosome-scale assembly of the lablab genome - A model for inclusive orphan crop genomics

**DOI:** 10.1101/2022.05.08.491073

**Authors:** Isaac Njaci, Bernice Waweru, Nadia Kamal, Meki Shehabu Muktar, David Fisher, Heidrun Gundlach, Collins Muli, Lucy Muthui, Mary Maranga, Davies Kiambi, Brigitte L Maass, Peter MF Emmrich, Jean-Baka Domelevo Entfellner, Manuel Spannagl, Mark A Chapman, Oluwaseyi Shorinola, Chris S Jones

## Abstract

Orphan crops (also described as underutilised and neglected crops) hold the key to diversified and climate-resilient food systems. After decades of neglect, the genome sequencing of orphan crops is gathering pace, providing the foundations for their accelerated domestication and improvement. Recent attention has however turned to the gross under-representation of researchers in Africa in the genome sequencing efforts of their indigenous orphan crops. Here we report a radically inclusive approach to orphan crop genomics using the case of *Lablab purpureus* (L.) Sweet (syn. *Dolichos lablab*, or hyacinth bean) – a legume native to Africa and cultivated throughout the tropics for food and forage. Our Africa-led South-North plant genome collaboration produced a high-quality chromosome-scale assembly of the lablab genome – the first chromosome-scale plant genome assembly locally sequenced in Africa. We also re-sequenced cultivated and wild accessions of lablab from Africa confirming two domestication events and examined the genetic diversity in lablab germplasm conserved in Africa. Our approach provides a valuable resource for lablab improvement and also presents a model that could be explored by other researchers sequencing indigenous crops particularly from Low and middle income countries (LMIC).

## Introduction

Three major crops currently provide more than 40% of global calorie intake^1^. This over-dependence on a few staple crops increases the vulnerability of global food systems to environmental and social instabilities. One promising strategy to diversify food systems is to improve the productivity and adoption of climate-resilient but underutilised orphan crops through genome-assisted breeding^2^.

Genome-assisted breeding offers hope of a new green revolution by helping to uncover and unlock novel genetic variation for crop improvement. Over the last 20 years, the genomes of 135 domesticated crops have been sequenced and assembled^3^, including those of orphan crops^2^. However, it has recently been acknowledged that researchers from Africa are grossly underrepresented in the genome sequencing efforts of their indigenous orphan crops^3,4^. None of the assemblies of native African plant species released till date were sequenced in Africa. The acute lack of sequencing facilities and high-performance computing infrastructures as well as bioinformatics capacity to handle big genome data, has meant that researchers in Africa have historically taken the back seat in most genome sequencing efforts^5^.

Here we present a model to overcome this under-representation through an inclusive orphan crop genomics approach. We applied an Africa-led, internationally collaborative approach to the genome sequencing of lablab (*Lablab purpureus* L. Sweet) - a tropical legume native to Africa (Figure 1A). Lablab is remarkably drought-resilient and thrives in a diverse range of environments, as such it is widely cultivated throughout the tropical and subtropical regions of Africa and Asia^6^. Lablab is a versatile multipurpose crop that contributes towards food, feed, nutritional and economic security, and is also rich in bioactive compounds with pharmacological potential, including against SARS-Cov2^7–10^. Climate change is driving researchers to investigate crops like lablab for its outstanding drought tolerance^11^.

**Figure 1:**
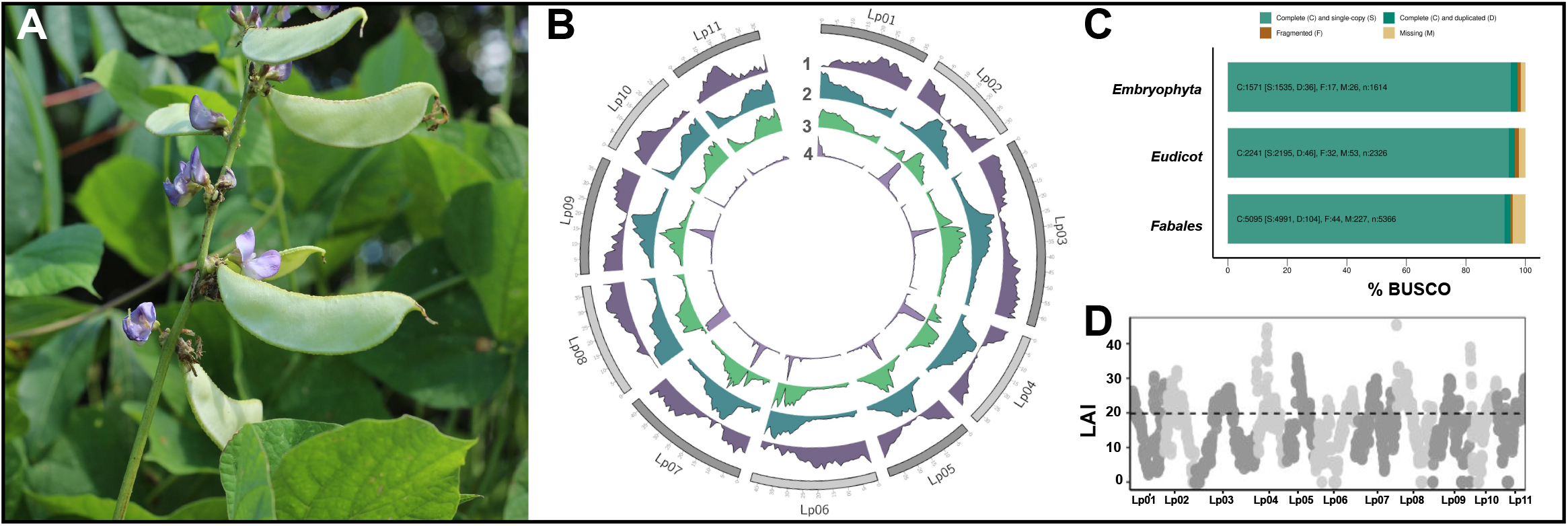
Genome Assembly of Lablab. **a**, Lablab purpureus plant showing flowers, leaves and pods; **b**, Gene and repeat landscape of the lablab genome. The tracks from the outer to the inner track show 1) Gene density, 2) Repeat density, 3) LTR-RT density, 4) Tandem repeat density. **c**, LAI index of the 11 lablab chromosomes; **d**, BUSCO scores of the Lablab purpureus genome annotation using the embryophyta, eudicots and and fabales reference lineages.

Our Africa-led genome collaboration produced a chromosome-scale assembly of lablab – the first chromosome-scale plant genome assembly sequenced in Africa. We also discuss the main features and benefits from our inclusive approach, and suggest this can serve as a roadmap for future genomic investigations of indigenous African crops.

## Results

### Genome sequencing

High acquisition and maintenance cost of sequencing platforms is a major limiting factor to genomics research in Africa. To circumvent this limitation, we used the portable and low-cost Oxford Nanopore Technology (ONT) MinION platform for in-country sequencing of the genome of lablab (cv. Highworth). We generated 4.7 M reads with a mean read length of 6.1 Kbp (Table S1). This amounted to 28.4 Gbp of sequences and 67x coverage of the lablab genome based on a previously estimated genome size of 423 Mbp^12^. The reads were initially assembled into 2,260 contigs with an N50 of 11.0 Mbp. The assembly was polished for error correction using ~380x of publicly available Illumina short reads (NCBI Bioproject PRJNA474418).

Using high-throughput Chromosome Conformation Capture (Hi-C), we clustered and oriented the contigs into 11 pseudomolecules covering 417.8 Mbp (98.6% of the estimated genome size) with an N50 of 38.1 Mbp (Figure 1B, Table S2, Supplemental methods). Our chromosome-scale assembly of the lablab genome has 61-fold improvement in contiguity compared to the previously published short read assembly^12^. For consistency with published legume genome sequences, we assigned chromosome names based on syntenic relationship with *Phaseolus vulgaris* (common bean^13^) and *Vigna unguiculata* (cowpea^14^) (Figure S1, Supplemental methods).

### Genome annotation and gene family analyses

To annotate the genome, we established an international collaboration comprising locally trained African researchers (see discussion) and international partners with established genome annotation pipelines. We used an automated pipeline based on protein homology, transcript evidence and *ab initio* predictions to identify protein coding genes in the lablab genome. This resulted in a total of 30,922 gene models (79,512 transcripts). A subset of 24,972 of these gene models show no homology to transposable elements (TEs) and can be confidently considered as high quality proteincoding non-TE gene models (Figure 1B, Table S3). BUSCO scores of the non-TE gene models were 97.3%, 96.4%, and 94.9% against the universal single copy genes from the embryophyta, eudicots, and fabales lineages, respectively, suggesting a high level of completeness of the gene space (Figure 1C). The number of non-TE protein-coding genes identified in our study is 19.2% greater than in the previous short-read assembly^12^. A functional description could be assigned to 28,927 (93.3%) of the genes.

A total of 168,174 TE sequences, occupying 28.1% of the genome, were identified in the lablab genome (Figure 1B). Of these, 89.6% were classified into 13 superfamilies and 2,353 known families (Table S4, Figure S2). Long Terminal Repeat - RetroTransposons (LTR-RTs) were the most abundant TEs, with 85,149 sequences occupying 83 Mb (19.9%) of the genome (Figure 1B). Copia were the most abundant LTR-RT superfamily, occupying 13.2% of the genome compared to gypsy elements that occupied only 4.7%. We also report an average LTR Assembly Index (LAI) of 19.8 (Figure 1D). DNA transposons were smaller in number and size relative to LTR-RTs, and were distributed more evenly across the chromosomes (Table S4, Figure S2).

A further 100,741 repetitive sequences were identified but could not be classified as TEs. Combining the annotated TEs and unclassified repeats reveals an overall repeat content of 43.4% of the genome. We also identified 142,302 tandem repeats (TRs) covering 43 Mb (11.2%) of the genome (Figure 1B, Table S5). Most of these were minisatellites (10-99 bp), while satellite repeats (>100 bp) make up the largest total proportion of TRs in the genome (7.4% of the genome; Table S5). Both the tandem and unclassified repeats were found to concentrate within a distinct, overlapping cluster at the point of peak repeat density on each chromosome, indicating that they are likely centromeric repeats (Figure S2A).

Gene family analysis and comparison to other legumes (*P. vulgaris*, *V. angularis*, *Cajanus cajan*, *Medicago truncatula*), and using *Arabidopsis thaliana* as an outgroup, placed 24,397 (97.7%) of the 24,972 non-TE lablab genes into orthogroups. Comparison of the five legumes (Figure 2A) revealed 14,047 orthogroups in common, and identified 417 (1.7%) lablab genes in 119 species-specific orthogroups that were absent from the other four legumes. These lablab-unique gene families were enriched for fatty acid biosynthesis, arabinose metabolism gene ontology (GO) classifiers while several were involved in pollen-pistil interactions and general plant development (Table S6). Using the phylogenetic relationships between the species, 448 gene families were significantly expanded in lablab compared to other legumes and *Arabidopsis*, while 899 were contracted (Figure 2B). Expanded gene families were enriched for lignin and pectin metabolism, photosynthesis among others (Table S7; Figure 2C).

**Figure 2:**
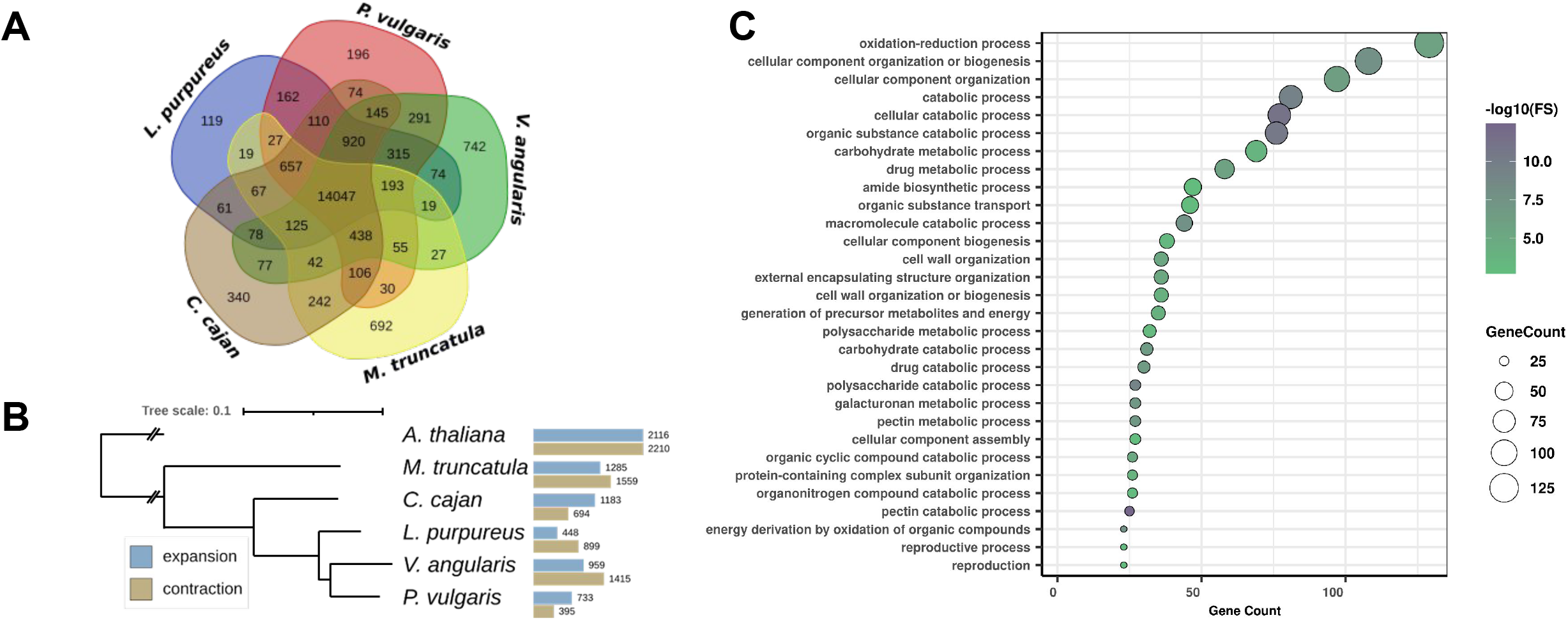
Gene family evolution and expansion in *Lablab purpureus*. **a**, Venn diagram of the number of gene families common among and unique to *Lablab purpureus, Phaseolus vulgaris, Vigna angularis, Medicago truncatula*, and *Cajanus cajan*. **b**, Cladogram of the analysed species showing the number of expanded and contracted gene families in each. Figure constructed with iTol^89^. **c**, Gene ontology terms enriched in the set of expanded gene families in *Lablab purpureus*.

### Evidence for two domestications of lablab

Understanding the transition from wild species to domesticated crop can provide insight into the location of domestication, the strength of genetic bottleneck (and identification of wild alleles not present in the domesticated gene pool) and can lead to identifying candidate genes underlying domestication traits. Previous work has suggested that lablab domestication occurred at least twice, separately in the two-seeded and four-seeded gene pools^15,16^. Using our chromosome-scale assembly as a reference, we examined whether this is indeed the case by resequencing a panel of two-seeded and four-seeded wild (ssp. *uncinatus*) and domesticated (ssp. *purpureus*) lablab accessions (Table S8). We also gathered publicly available short read data for cv. Highworth and nine species from three related genera (*Vigna*, *Phaseolus* and *Macrotyloma*, Table S8) as outgroups to determine the phylogenetic position of lablab. All lablab samples had a >95% mapping against the lablab reference genome at a depth of 7.0 - 11.2x while the related genera had considerably lower mapping of 30 - 54% at a depth of 3.5 - 10.9x; Table S8). A total of 39,907,704 SNPs were identified across all 22 samples and 15,428,858 across the 13 lablab samples.

A filtered SNP data set of 67,259 SNPs (see Methods) was used for phylogenetic and diversity analyses. Neighbor Joining phylogenetic analysis rooted with two *Macrotyloma* samples revealed that all lablab samples formed one group separate from the *Vigna* and *Phaseolus* samples which are each reciprocally monophyletic. A clear division between the two- and four-seeded lablab samples could be observed (100% bootstrap support) with wild and domesticated samples found in both groups (Figure 3). Our study thus confirms the previous hypotheses of two origins of domesticated lablab. Genetic diversity (π per 100 Kb window) within each gene pool was relatively low and significantly greater (unpaired T-test, t = 8.2415, df = 2651, P < 0.0001) in the two-seeded group (5.79 × 10^-6^ (+/− 2.51 × 10^-6^ [SD]) than the four-seeded group (5.04 × 10^-6^ (+/− 2.15 × 10^-6^ [SD]). Divergence between the two- and four-seeded gene pools was high (mean Fst per 100kb window = 0.43 +/− 0.32 [SD]) which could suggest that these gene pools should be taxonomically re-evaluated as separate species.

**Figure 3:**
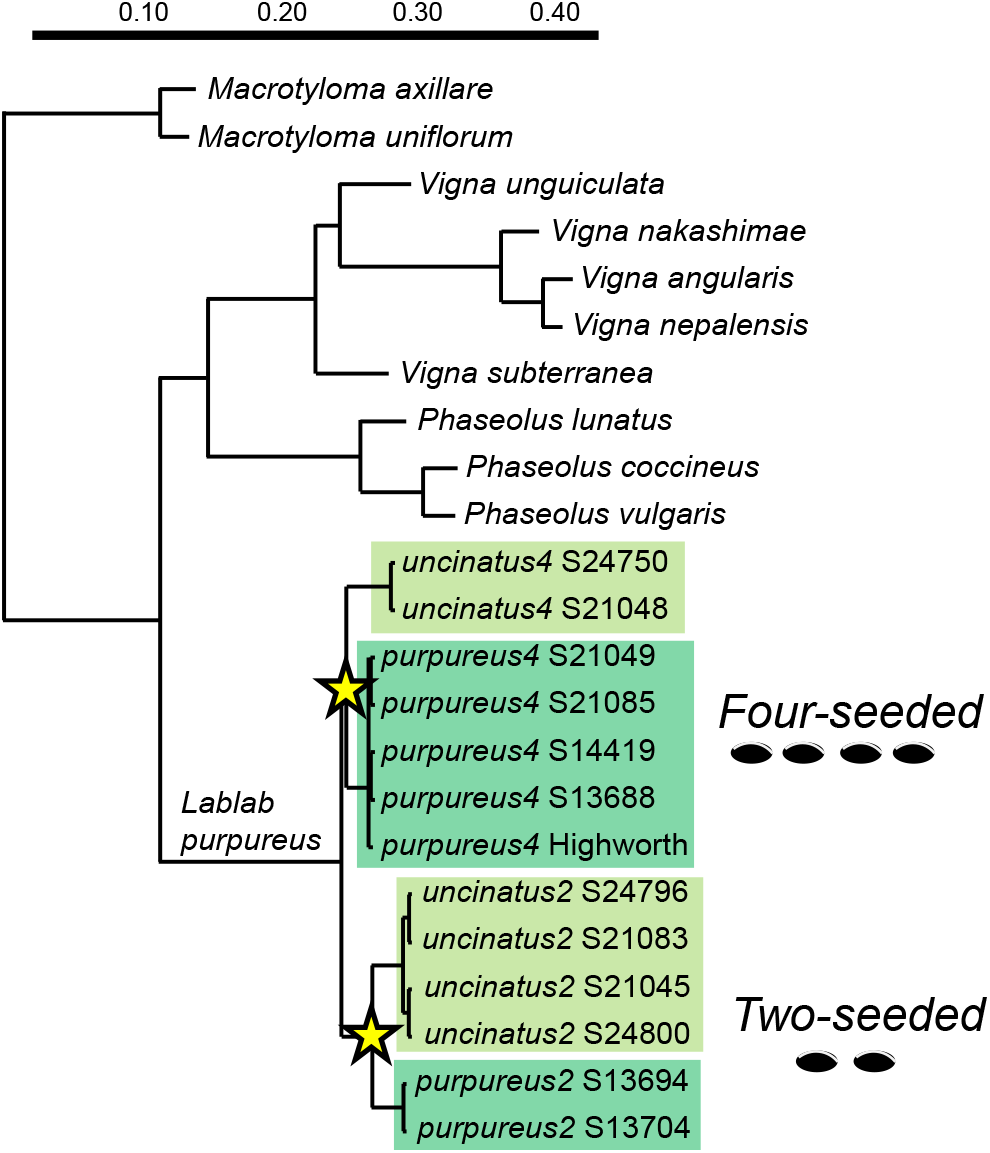
Phylogenetics of lablab and related legumes. Neighbor Joining phylogenetic relationships among lablab samples (2-seeded and 4-seeded purpureus (domesticated) and uncinatus (wild) subspecies) and other related legumes (see Table S8 for details). Tree is rooted on Macrotyloma. All nodes received full (100%) bootstrap support. Asterisks indicate the two domestication events.

### Genetic diversity in a global lablab collection

To assess within and between accession diversity in the global lablab gene pool, we genotyped 1,860 individuals from 166 lablab accessions using DArTseq genotyping-by-sequencing (GBS) (Table S9). We identified 41,718 genome-wide SNP and 73,211 SilicoDArT markers, of which 91% and 57% mapped onto the lablab genome, respectively (Figure S3). The two-seeded and wild samples mapped with a significant amount of missing data (due to the high genetic divergence described above), therefore we excluded these and report only results for the widespread four-seeded cultivated group. In addition, only individuals that were considered true-to-type or progeny (see Supplementary Information) were included. This resulted in 1,462 individuals from 138 accessions being retained for the final analysis.

Using a subset of 2,460 quality-filtered genome-wide SNPs (see Methods) for STRUCTURE^17^ analysis, we identified four populations (cluster I - IV) in the lablab germplasm collection (Figure 4A).

**Figure 4:**
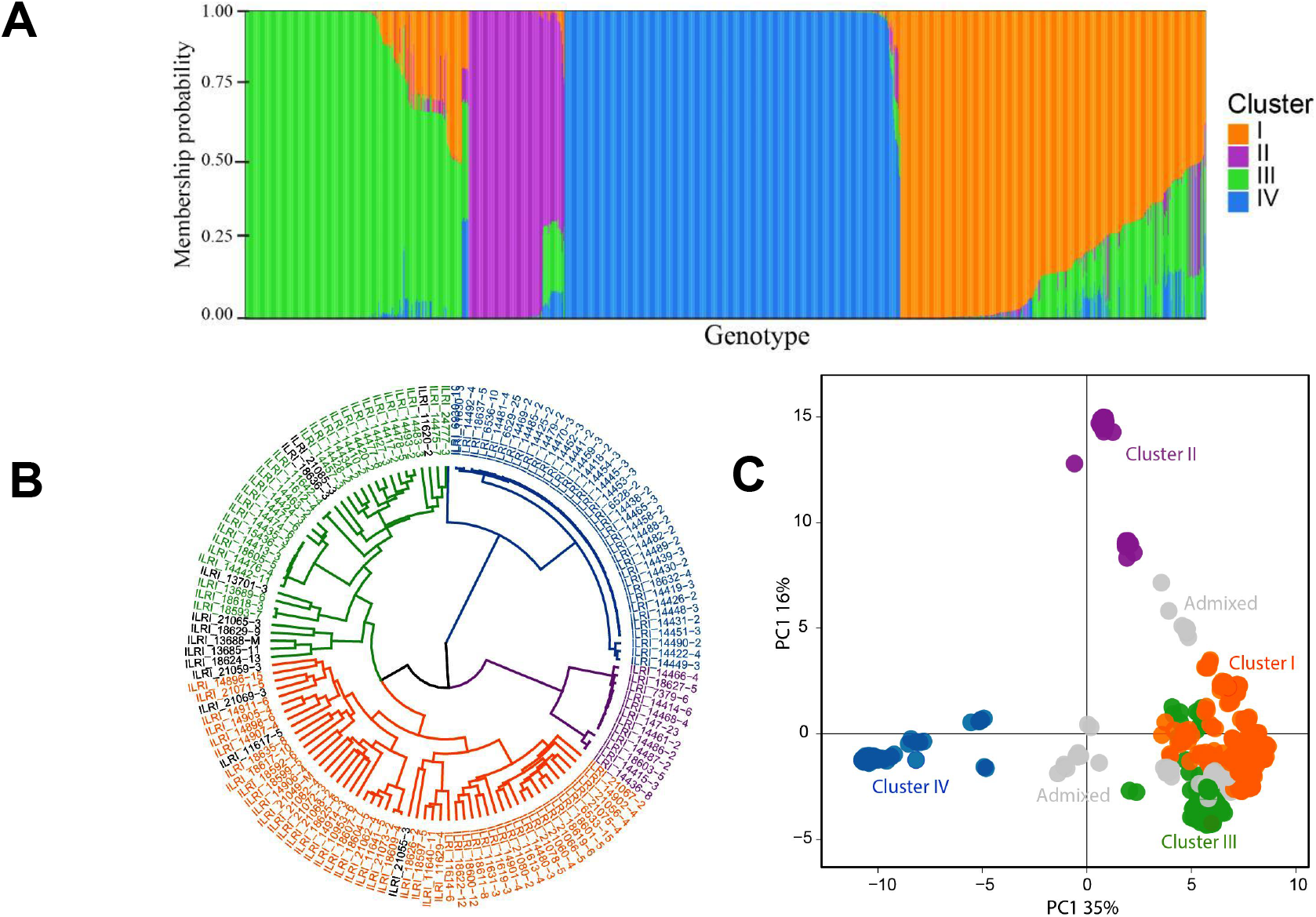
Clusters and subclusters of the lablab accessions used in the diversity study. **a**, Bar plots based on the admixture model in STRUCTURE for K = 4 (Membership of individual accessions to each subgroup is given in Table S16). **b**, Clusters detected by hierarchical clustering. **c**, Clusters detected by PCA. The colours in **b** and **c** are according to the STRUCTURE analysis in **a**.

Similar clustering and population stratification were detected by hierarchical clustering and PCA (Figure 4B and C). The clustering shows some correspondence with the geographical origin of the genotypes. Accessions in cluster I were mainly from outside Africa and included all the accessions of ssp. *bengalensis*, which has long, relatively narrow pods with up to seven seeds and a particular seed arrangment in the pod. More than 85% of the accessions in clusters II, III and IV are from Africa or were originally collected by the Grassland Research Station in Kitale (Kenya, but most have uncertain origin, Table S9).

The pairwise Fst values among the four clusters varied from 0.31 between clusters I and III to 0.91 between clusters II and IV (Table S10). Analysis of molecular variance (AMOVA) further showed presence of higher genetic variation between the four clusters (62.44%) than within the clusters (37.56%) (Table S11). Within group genetic distance between accessions, Nei’s D^18^, was lowest within cluster IV (mean D = 0.003) and highest for cluster I (mean D = 0.164; Table S12). Mean Nei’s D between progenies of the 41 accessions with ≥2 progenies per accession ranged from 0.0015 to 0.1516 indicating that within accession genetic diversity is generally low, as expected for a predominantly self-pollinating species such as lablab^19^.

We found that the population clusters often differed in their mean phenotypes based on historical data describing phenology and morpho-agronomic traits^20^. Twelve of 13 quantitative traits (Figure S4; Table S13) and five of eight qualitative characters (Figure S5; Table S14) differed among the four clusters despite a certain level of phenotypic variation within every cluster. Cluster I accessions are phenotypically variable, containing early-flowering, short plants and includes the only three erect accessions and all ssp. *bengalensis* in a sub-cluster. Plants had four to six relatively large seeds per pod. Cluster II contains the earliest, only colored-flowering accessions, with high flowering node density, and most producing up to four black, mottled seeds per pod. Plants were rather short and had the smallest leaves. Cluster III also includes diverse phenotypes; overall plants were relatively tall, broad, leafy and intermediate to late-flowering with the largest leaves and shortest pods with up to four rather small seeds. Cluster IV comprises the most homogeneous phenotypes; it had the latest, only white-flowering accessions and plants were rather tall, broad and leafy with long flower peduncles, a high number of flowering nodes and four relatively small tan-colored seeds per pod.

## Discussion

Africa has a rich plant biodiversity that includes 45,000 species^21^, most of which are under-studied and under-utilised. To fully explore these genetic resources, it is important to develop inclusive research models that enable and empower local researchers to study these species under a resource-limited research setting. Our work describes an inclusive African-led effort to produce high-quality genome resources for a climate-resilient and multipurpose native African orphan crop - lablab. Our chromosome-scale reference assembly of lablab improves on the previous assembly in several ways and also highlights some interesting features about lablab’s genome, domestication and diversity.

With the use of long-reads and Hi-C scaffolding, we achieved 61-fold improvement in contiguity, and identified a further 34 Mbp of repetitive sequences and 19.2% more gene content compared to the short-read based assembly^12^. In addition, the high average LTR Assembly Index (LAI)^22^ (19.8; Figure 1C), comparable to the LAI of a PacBio-based assembly of common bean^23^, indicates a high-level of completeness of the repeat space in our assembly. As has been found in other legumes, LTR-RTs were the predominant TE class in our lablab assembly^13,14,24^. In contrast to findings from lablab’s close relatives, however, we found copia LTR-RTs to be more abundant than gypsy LTR-RTs. It is uncommon to see a greater abundance of copia LTR-RTs when compared to gypsy LTR-RTs in plant genomes^25,26^, and although the biological significance of elevated copia abundances remains to be seen, further genome sequencing will determine whether this finding is indeed a distinguishing feature of lablab.

Lablab has a smaller genome size than other sequenced legumes and also has a smaller number of species-specific orthogroups. Nevertheless, the orthogroup analysis identified several GO categories enriched in the lablab-specific orthogroups; of particular interest are those involved in fatty acid metabolism, which could underlie seed oil content and composition. In addition arabinose metabolism genes were enriched in the lablab-unique genes and several other cell wall-related GO terms (specifically related to pectin and lignin) in the orthogroups expanded in lablab. Cell wall modification could be related to protection from pathogens^27^ or drought tolerance^28^.

A dual origin of domesticated lablab was confirmed, with the localised (to Ethiopia) two-seeded and the widespread four-seeded types being genetically distinct and domestication events occurring in both of these groups. This therefore adds lablab to the relatively ‘exclusive’ list of crops with more than one origin, which includes common bean^13^, lychee^29^, Tartary buckwheat^30^ and, potentially, rice^31^ and barley^32^. Data on reproductive isolation between the gene pools is unclear, and crosses are only known between four-seeded samples^33–35^, thus any taxonomic reassessment (first suggested by Maass et al. 2005^15^) should begin with assessing reproductive compatibility between the gene pools.

Importantly, our project provides a model for increasing the representation of local researchers in the sequencing of their indigenous crops. Recent studies and commentaries have highlighted the disconnect between the species origin and the location of the institutions leading their sequencing^3–5^. This is particularly true for Africa, where none of its sequenced indigenous crops were sequenced on the continent^4^. We surveyed 31 publications describing the genome sequencing of 24 indigenous African crops. More than 85% of these publications do not have first or corresponding authors with affiliations in Africa and 42% do not have any authors with an African affiliation (Table S15). Our project breaks this trend because sequencing and coordination efforts were done or led from within Africa, while still recruiting international partners where complementary expertise was beneficial to the project. Thus we encourage contribution of the international community in African orphan crop genomics while supporting more active involvement from local researchers.

Three main features characterised our inclusive genome collaboration model - access to low-cost portable sequencing, in-depth capacity building and equitable international collaboration. The high acquisition and maintenance costs of genome sequencing technologies has historically limited the participation of researchers working in LMIC in genome collaborations. Low-cost and portable sequencing platforms such as the ONT MinION, are now making long-read sequencing accessible to researchers in LMIC, thus “democratising” genome sequencing. We spent less than $4,000 to procure the MinION sequencer and kits (ONT starter pack and extra flow cells) used to sequence our lablab genome. This low cost is partly due to the small genome size of lablab, but it nonetheless demonstrates how accessible modern portable sequencing platforms can be for researchers in resource-limited research settings. Despite these low costs, there are, however, still logistical challenges to overcome in getting needed reagents to local labs.

Secondly, our project benefited from efforts to build in-depth bioinformatics skills in Africa^5^. Four of the African authors in our study, including two of the first authors, benefited from a residential 8-month bioinformatics training in Africa. We posit that such in-country and long-term training, as opposed to short training, are more effective in developing the high-competence bioinformatics skills that the continent needs. Once trained, these researchers will feel empowered to participate or lead genomic projects, and importantly use such projects as opportunities to train many more researchers, thus creating a continuous stream of human resources equipped to explore the rich genetic resources on the continent.

Lastly, establishing an international collaboration helped us to take advantage of existing expertise and already developed pipelines for genome analyses. With over 20 years of plant genome sequencing, the global plant science community have developed tools, pipelines and protocols for plant genome analyses. This means African researchers do not have to ‘reinvent the wheel’ for orphan crop genomics, but instead can form strategic collaborations to access needed expertise and networks. To fully benefit from big-data and a suite of readily-available genomic tools, it is also vital that African institutions are supported to build or access physical or cloud computing infrastructure for high-throughput data analytics. This will also ensure that genomic data produced on the continent are locally managed and made readily accessible to local researchers and the global community.

Our lablab genome assembly and collaboration provides a roadmap for improving agronomic, yield and nutritional traits in other African orphan crops. Given the Africa-centred and inclusive nature of our work, this could be used as a model by individual labs and multinational genome consortia including the Africa Biogenome Initiative^3^ to generate high-quality genomic resources for many indigenous species across the continent.

## Methods

### Reference genome DNA extraction and sequencing

*L. purpureus* (L.) Sweet cv. Highworth^36^ seeds were germinated in a petri dish on filter papers moistened with tap water. The sprouted seedlings were transferred to soil and allowed to grow for one month in the greenhouse facility at the International Livestock Research Institute (ILRI, Kenya). Two grams of young trifoliate leaves were harvested, flash frozen in liquid nitrogen and stored at −80°C. The leaves were ground in liquid nitrogen using a pestle and mortar and High Molecular Weight (HMW) DNA extracted with Carlson lysis buffer (100 mM Tris-HCl, pH 9.5, 2% CTAB, 1.4 M NaCl, 1% PEG 8000, 20 mM EDTA) followed by purification using the Qiagen Genomic-tip 100/G based on the Oxford Nanopore Technologies (ONT) HMW plant DNA extraction protocol. The library was prepared following the ONT SQK-LSK109 ligation sequencing kit protocol. A total of 1 μg of genomic DNA was repaired and 3’-adenylated with the NEBNext FFPE DNA Repair Mix and the NEBNext^®^ Ultra^™^ II End Repair/dA-Tailing Module and sequencing adapters ligated using the NEBNext Quick Ligation Module (NEB). After library purification with AMPure XP beads, sequencing was conducted at ILRI (Kenya) using the R9.4.1 flow cells on a MinION sequencer platform.

### Genome Assembly

Guppy basecaller (v4.1.1)^37^ was used for base calling the reads using the high accuracy basecalling model and the resulting fastq files were used for genome assembly. Flye *de novo* long reads assembler^38^ (ver 2.7.1) was used for the assembly with the default parameters. The draft assembly was polished with lablab Illumina shorts reads^12^ using HyPo hybrid polisher^39^. The draft genome assembly quality was assessed using QUAST^40^ and its completeness evaluated using BUSCO (ver. 4.0.6)^41^. The Hi-C library for genome scaffolding was prepared, sequenced and assembled by phase genomics, USA (Supplemental Information).

### Gene Annotation

Protein sequences from five closely related species (*P. vulgaris*, *V. angularis*, *C. cajan*, and *M. truncatula*) as well as *Arabidopsis thaliana* were used as protein homology evidence. RNAseq data from *Lablab purpureus* cv. Highworth leaves, stem, sepals, and petals^12^ was used in *de novo* transcript assembly with Trinity^42^ (ver 2.8.5) and provided as transcript evidence. The Funannotate pipeline^43^ (ver 1.8.7) was used for gene prediction using RNA-Seq reads, *de novo* assembled transcripts and soft-masked genome as input to generate an initial set of gene models using PASA^44^ (ver 2.4.1). Next, the gene models and protein homology evidence were used to train Augustus^45^ (ver 3.3.3), SNAP^46^ (ver 2006-07-28) and Glimmerhmm^47^ (ver 3.0.4) *ab initio* gene predictors and predicted genes passed to Evidence modeller^48^ (ver 1.1.1) with various weights for integration. tRNAscan-SE^49^ (ver 2.0.9) was used to predict non-overlapping tRNAs. Transcript evidence was then used to correct, improve and update the predicted gene models and refine the 5’- and 3’-untranslated regions (UTRs).

The plant.annot pipeline (github.com/PGSB-HMGU/plant.annot) was also used for the prediction of protein coding genes and incorporated homology information and transcript evidence as well. In the evidence-based step, RNA-Seq data from cv. Highworth leaf, stem, sepal and petal^12^ was used for the genome-guided prediction of gene structures. HISAT2^50^ (version 2.1.0, parameter –dta) was used to map RNA-Seq data to the reference genome and the transcripts assembled with Stringtie^51^ (version 1.2.3, parameters -m 150 -t -f 0.3). For the homology-based step, homologous proteins from the closely related species were mapped to the reference genome using the splice-aware mapper GenomeThreader^52^ (version 1.7.1, parameters: -startcodon -finalstopcodon -species medicago - gcmincoverage 70 -prseedlength 7 -prhdist 4). Transdecoder^53^ (version 3.0.0) was used to predict protein sequences and to identify potential open reading frames. The predicted protein sequences were compared to a protein reference database (UniProt Magnoliophyta, reviewed/Swiss-Prot) using BLASTP^54^ (-max_target_seqs 1 -evalue 1e^-05^). Conserved protein family domains for all proteins were identified with hmmscan^55^ version 3.1b2. Transdecoder-predict was run on the BLAST and hmmscan results and the best translation per transcript was selected. Results from the homology and transcriptbased gene prediction approaches were combined and redundant protein sequences were removed.

The results from both the funannotate and plant.annot pipelines were combined and redundant protein sequences as well as non-coding genes removed. The functional annotation of transcripts as well as the assignment of Pfam^56^- and InterPro^57^-domains, and GO^58,59^ terms, were performed using AHRD (Automatic assignment of Human Readable Descriptions, https://github.com/groupschoof/AHRD; version 3.3.3). AHRD assesses homology information to other known proteins using BLASTP searches against Swiss-Prot, The Arabidopsis Information Resource (TAIR), and TrEMBL. The functional annotations are defined using the homology information and the domain search results from InterProScan and Gene Ontology terms. In order to distinguish transposon related genes from other genes, the functional annotation was used to tag TE-related genes in the genome annotation file. BUSCO^41^ v5.2.2 was used to assess the completeness of the genome annotation, with sets of universal single copy gene orthologs from embryophyta, fabales, and eudicots odb10 lineages^41^.

### Repeat Annotation

Repeat annotations for transposable elements (TE) and tandem repeats were conducted independently. For TE annotation, a novel Lablab TE library was constructed using the Extensive de novo TE Annotator (EDTA v1.9.7) pipeline^60^. EDTA incorporates both structure and homology-based detection programs to annotate the predominant TE classes found in plant genomes. EDTA utilises LTRharvest^61^, LTR_FINDER^62^, LTR_retriever^63^, TIR-Learner^64^, HelitronScanner^65^, RepeatModeler2^66^ and RepeatMasker^67^ for identification of novel TE sequences. The outputs of each module are then combined and filtered to compile a comprehensive, non-redundant TE library. EDTA’s inbuilt whole genome annotation function was then used to produce a non-overlapping TE annotation for lablab using the TE library as input. Further calculation of metrics and data visualisation were carried out in R^68^ using the tidyverse suite^69^ of packages.

Tandem repeats were identified with TandemRepeatFinder^70^ under default parameters and subjected to an overlap removal by prioritising higher scores. Higher scoring matches were assigned first. Lower scoring hits at overlapping positions were either shortened or removed. Removal was triggered if the lower scoring hits were contained to ≥ 90% in the overlap or if less than 50 bp of rest length remained.

### Gene family and expansion analysis

Gene families were identified using a genome-wide phylogenetic comparison of the lablab protein sequences and four other legumes. This comprised *P. vulgaris* (PhaVulg1_0)*, V. angularis* (Vigan1.1), *C. cajan* (V1.0), and *M. truncatula* (MtrunA17r5). Orthofinder^71^ (Version 2.4) was used to identify orthologs and co-orthologs between these species and to group them into gene families. *Arabidopsis thaliana* (Araport 11) was used as an outgroup. The longest transcript was selected for genes with multiple splice variants.

In order to analyse gene family expansion and contraction in lablab, the gene family file produced by Orthofinder was further analysed with CAFE5^72^. An ultrametric tree was built with Orthofinder (r=160) and CAFE5 ^72^ was run with -k 3. Enrichment analysis using a fisher’s exact test (padj ≤ 0.05) of significantly (p-value of gene family sizes^73^ ≤ 0.05) expanded gene families was performed with TopGO^74^.

### Resequencing and Phylogenetic Analyses

Lablab seeds (obtained from ILRI) for the resequencing were germinated in a 1:1 mixture of vermiculite and Levingtons’s M2+S compost in a greenhouse (22°C and 16 hour day) at the University of Southampton. Young leaf tissue was harvested from one-month old seedlings and snap frozen in liquid nitrogen. DNA was extracted from leaf tissue using a CTAB-based protocol^75^ with minor modifications. In total, 12 samples from two and four-seeded wild and domesticated lablab accessions were sequenced using 2 × 150 bp PE sequencing on an Illumina platform at Novogene (Cambridge, UK) (Table S6). Short read data from lablab cv. Highworth^12^, three *Phaseolus*, four *Vigna*, and two *Macrotyloma* species were downloaded from the NCBI Sequence Read Archive (Table S8). A maximum of 100 M read pairs were downloaded.

The reads were trimmed using Trimmomatic^76^ (ver 0.32) with the parameters; ILLUMINACLIP:TruSeq3-PE-2.fa:2:30:10, LEADING:5, TRAILING:5, SLIDINGWINDOW:4:15, MINLEN:72. Between 21.9 and 97.1 M reads remained after trimming. The trimmed reads were mapped to the chromosome-scale lablab assembly (excluding unmapped contigs) using Bowtie2^77^ (ver 2.2.3) and *--very-sensitive-local* settings. SAMtools^78^ (ver 1.1) was used to convert .sam to .bam files which were then sorted, and duplicated reads were removed using the Picard toolkit^79^ (ver 2.8.3, VALIDATION_STRINGENCY=LENIENT). Depth was estimated using SAMtools^78^ (Table S6). Using mpileup from bcftools^80^ (ver 1.6.0), the individual sorted bam files were combined into a multi-sample VCF using the settings *-Q 13* and *-q 10* and variant detection was performed with “bcftools call”. Variants were subsequently filtered using “bcftools filter”, *-i’QUAL>20 & DP>6’*. The proportion of missing data per individual was calculated using vcftools^81^ (ver 0.1.14; Table S6). Finally, vcftools was used to trim the filtered VCF, removing SNPs that were missing in more than two samples and those with a minor allele frequency of <5%. Finally, only SNPs that were at least 2 Kbp apart were included. The final file contained 67,259 SNPs. VCF2Dis (github.com/BGI-shenzhen/VCF2Dis/; ver 1.36) was used to create a distance matrix which was submitted to the FAST-ME server (atgc-montpellier.fr/fastme) to generate a NJ tree. A total of 1000 replicate matrices were generated in VCF2Dis and the phylip commands *“neighbor”* and *“consense”* were used to calculate bootstrap values. Genetic diversity for the two subpopulations and Fst between the subpopulations were calculated from the final VCF file using vcftools in 100kb windows.

### Population structure and diversity

A total of 1,860 seedlings from 166 *Lablab purpureus* accessions, that have been maintained at the ILRI forage genebank were grown from seed under screen house conditions at ILRI, Ethiopia. Genomic DNA was extracted from leaves using a DNeasy^®^ Plant Mini Kit (Qiagen Inc., Valencia, CA). The DNA samples were genotyped by the DArTseq genotyping platform at Diversity Arrays Technology, Canberra, Australia^82^. A subset of 2,460 robust SNP markers was filtered based on the marker’s minor allele frequency (MAF ≥ 2 %), missing values (NA ≤ 10 %), independence from each other (Linkage disequilibrium-LD ≤ 0.7), and their distribution across the genome.

A pairwise IBD (Identity-By-Descent) analysis was conducted using PLINK^83^ and contaminants excluded from the following analyses (see Supplemental Information) Genetic diversity was estimated using pairwise Nei’s genetic distance^18^. Population stratification was assessed using the Bayesian algorithm implemented in STRUCTURE^17^, in which the burn-in time and number of iterations were both set to 100,000 with 10 repetitions, testing the likelihood of 1-10 subpopulations in an admixture model with correlated allele frequencies. Using Structure Harvester^84^ the most likely number of subpopulations was determined by the Evanno ΔK method^85^. Accessions with less than 60% membership probability were considered admixed. Hierarchical clustering, principal component analysis (PCA), fixation index (Fst), and analysis of molecular variance (AMOVA) were conducted using the R-packages *Poppr*^86^, adegenet^87^, and APE^88^.

## Data availability

The lablab genome is available from NCBI BioProject (PRJNA824307) and at https://hpc.ilri.cgiar.org/~bngina/lablab_longread_sequencing_March_2022/. Raw sequencing reads for the resequencing are available from the NCBI SRA under project number PRJNA834808.

## Acknowledgements

This research was conducted as part of the CGIAR Research Program on Livestock, supported by CGIAR Fund Donors. O.S. was supported by the Royal Society FLAIR award (FLR_R1_191850), N.K., H.G. and M.S. were supported by the German Federal Ministry of Education and Research (De.NBI, FKZ 031A536). D.F. was supported by the SoCoBio DTP (grant number BB/T008768/1; BBSRC, UK) to carry out a PhD rotation project in the lab of MAC. DF and MAC acknowledge the use of the IRIDIS High Performance Computing Facility, and associated support services at the University of Southampton.

## Authors’ contributions

O.S., P.M.F.E., J.D.E., M.A.C., M.S. and C.S.J. conceived and planned the experiments. C.M., L.M. and O.S. performed DNA extraction and Nanopore Sequencing. I.S., B.W., M.M. and D.K. performed the genome assembly. N.K., B.W., M.S. and I.S. performed genome annotation. D.F. and H.G. annotated the transposable elements and tandem repeats. N.K., B.W. and O.S performed gene family analyses, M.A.C. analysed the re-sequencing data. M.S.M., C.S.J. and B.L.M. performed diversity analyses on global collection. I.N., B.W., N.K., M.S.M., D.F., M.A.C., O.S. and C.S.J. wrote the manuscript. All authors reviewed and approved the final manuscript.

## Materials & Correspondence.

Mark Chapman, Oluwaseyi Shorinola, Chris Jones.

## Supplementary Information

### Supplementary Methods

#### Hi-C Scaffolding

Chromatin conformation capture data was generated by Phase Genomics (Seattle, USA) using the Proximo Hi-C 2.0 Kit, which is a commercially available version of the Hi-C protocol. Following the manufacturer’s instructions for the kit, intact cells from two samples were crosslinked using a formaldehyde solution, digested using the DPNII restriction enzyme, end repaired with biotinylated nucleotides, and proximity ligated to create chimeric molecules composed of fragments from different regions of the genome that were physically proximal in vivo, but not necessarily genomically proximal. Continuing with the manufacturer’s protocol, molecules were pulled down with streptavidin beads and processed into an Illumina-compatible sequencing library. Sequencing was performed on an Illumina HiSeq, generating a total of 232,382,372 PE150 read pairs.

Reads were aligned to the draft assembly using BWA-MEM^90^ with the -5SP and -t 8 options specified, and all other options default. SAMBLASTER^91^ was used to flag PCR duplicates, which were later excluded from analysis. Alignments were then filtered with SAMtools^77^ using the -F 2304 filtering flag to remove non-primary and secondary alignments. Putative misjoined contigs were broken using Juicebox^92^ based on the Hi-C alignments. A total of 6 breaks in 6 contigs were introduced. The same alignment procedure was repeated from the beginning on the resulting corrected assembly.

Phase Genomics Proximo Hi-C genome scaffolding platform was used to create chromosome-scale scaffolds from the corrected assembly as described in Bickhart et al.^93^. As in the LACHESIS method^94^, this process computes a contact frequency matrix from the aligned Hi-C read pairs, normalized by the number of DPNII restriction sites (GATC) on each contig, and constructs scaffolds in such a way as to optimize expected contact frequency and other statistical patterns in Hi-C data. Approximately 20,000 separate Proximo runs were performed to optimize the number of scaffolds and scaffold construction in order to make the scaffolds as concordant with the observed Hi-C data as possible. This process resulted in a set of 11 chromosome-scale scaffolds containing 417 Mbp of sequence (98% of the corrected assembly) with a scaffold N50 of 38.1 Mbp.

#### Synteny-guided Chromosome naming

We adopted a naming scheme based on synteny with closely related legumes - *P. vulgaris* (common bean^13^) and *V. unguiculata* (cowpea^14^). For this, we downloaded protein sequence and GFF files of PacBio-based assembly of *P. vulgaris* (v2.1) and *V. unguiculata* (v1.2) from Phytozome^23^ and compared this separately to lablab proteins using BLASTP^54^ (settings: -max_target_seqs 1, -evalue 1e-10, -qcov_hsp_perc 70). MCScanX^95^ was used to process the individual BLAST output and to detect inter-species collinear blocks.

#### Filtering for true-to-type genotypes in global genebank collection

The lablab accessions used for evaluating global diversity in this study were acquired from different sources and conserved *ex situ* as seeds in the ILRI forage genebank, the earliest since 1982, with periodic monitoring for viability and regeneration for renewal of the seeds. These periodic genebank management practices involve risks to the genetic integrity of the accessions through pollen contamination, seed contamination, segregation, mislabeling, and other factors (e.g. as described in Chebotar et al., 2003 ^96^). Hence, it was necessary to ensure the genetic integrity of plants within accessions and avoid potential contaminants before the genetic diversity analysis. Using pairwise IBD (Identity-By-Descent) analysis, plants within accessions were classified into “true-to-type”, “progeny”, or “contaminant” based on a PI_HAT^83^ value of above 0.80, between 0.125 and 0.80, or less than 0.125, respectively. Six accessions with a single plant each were excluded from the analysis.

For nine accessions, all plants were unrelated to each other, and therefore considered “contaminants”. Out of the remaining 151 accessions, 85 were 100% true-to-type, indicating that there was no cross-pollination or seed mixing. Twenty-four accessions had a mixture of true-to-type and their progeny, indicating that some level of cross-pollination or segregation had taken place in this group. Another 24 accessions had a mixture of true-to-type and contaminants, and other 18 accessions had a mixture of the true-to-type, their progenies, and contaminants (Figure S6). After removing contaminants, a total of 1680 plants were retained from these 151 accessions for genetic diversity analysis. Of these, 1541 plants were true-to-type with 2 to 26 plants per accession, and 139 were progenies from 41 accessions (1 to 12 plants per accession).

#### Analysing historical lablab phenotype datasets

Phenotypic variation among the identified major molecular groups was assessed based on historical data summarised by Pengelly and Maass (2001)^20^ (127 accessions) and Wiedow (2001)^97^ (95 accessions), in which morpho-agronomic traits on lablab accessions were evaluated in field trials at Ziway site in Ethiopia, in 1998 and 2000, respectively. Seventeen accessions were analysed in both trials, hence we could determine whether traits varied across the seasons. Where variation was low (correlation between seasons was 80% or greater; 6 traits), data from the two trials were combined. For the remaining 15 traits, only the 1998 phenotype data on 75 accessions was used for the analysis of trait variation among the four genetic groups identified above. Analysis of variance (ANOVA) and Tukey’s multiple comparison test were employed to compare phenotypic variation of agro-morphological quantitative traits with significant p values (*P* < 0.01) among clusters identified by population structure analysis. A chi-square test was used for similar comparisons among clusters in qualitative traits.

## Supplementary Figures

**Figure S1:**
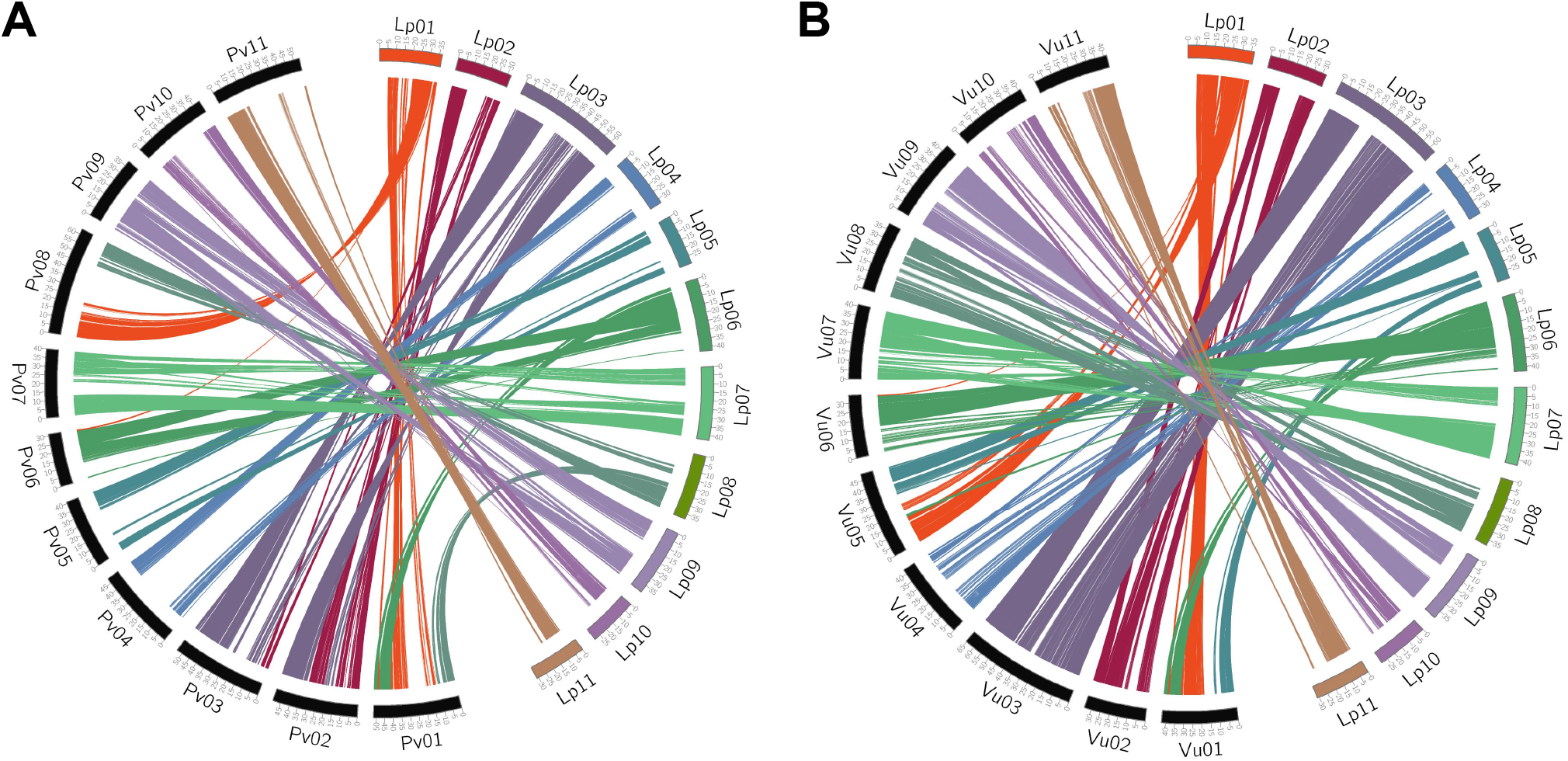
Chromosome-level synteny of *Lablab purpureus* with related species. *L. purpureus* chromosomes have been named according to synteny with *P. vulgaris* **(a)** and *V. unguiculata* **(b)** chromosomes.

**Figure S2:**
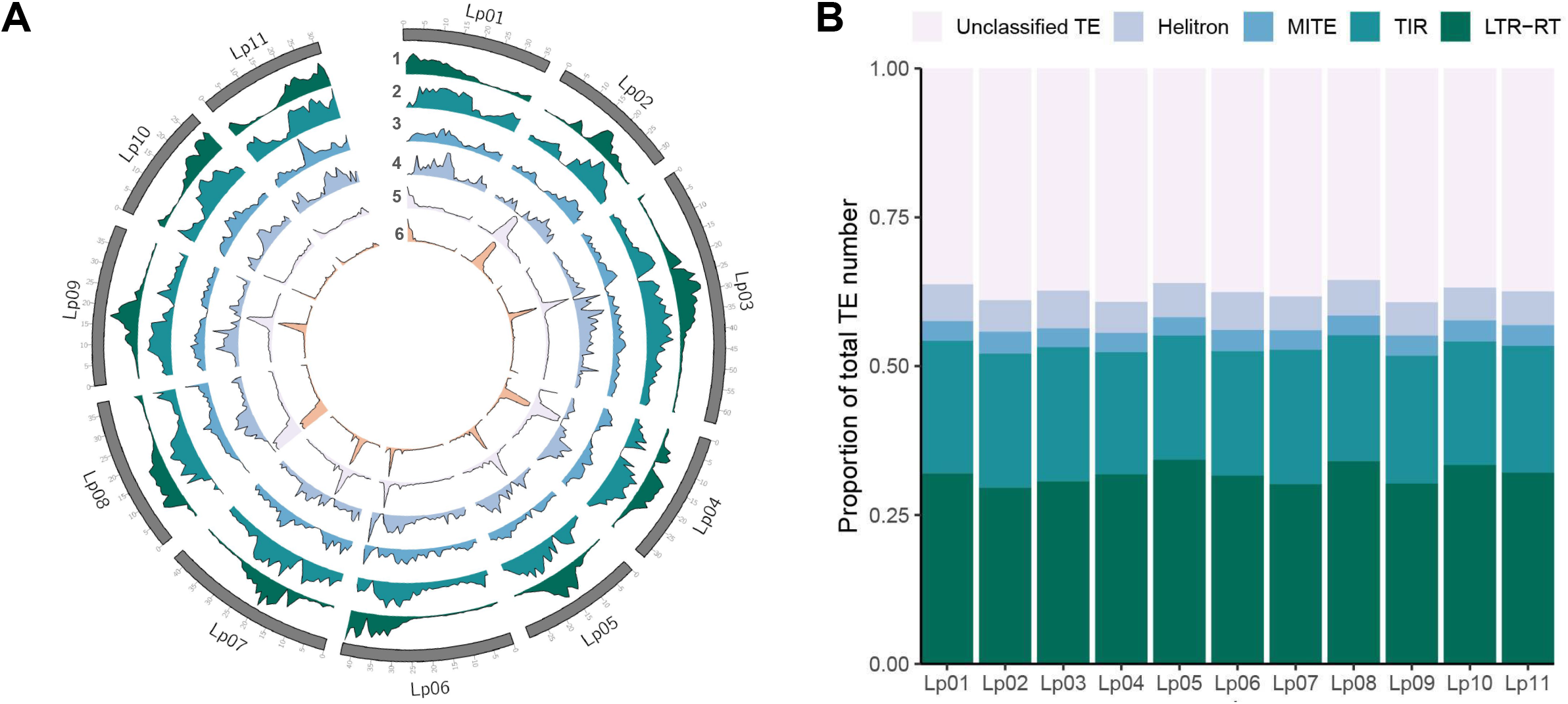
Chromosomal repeat content in *Lablab purpureus*. **(a)** Relative densities of repeat elements along each chromosome. 1) Long Terminal Repeat RetroTransposons (LTR-RT), 2) Tandem Inverted Repeats (TIR), 3) Miniature Inverted Transposable Elements (MITE), 4) Helitron 5) Unclassified repeats, 6) Tandem repeats **(b)** Proportional abundance of identified transposable element orders on each chromosome.

**Figure S3:**
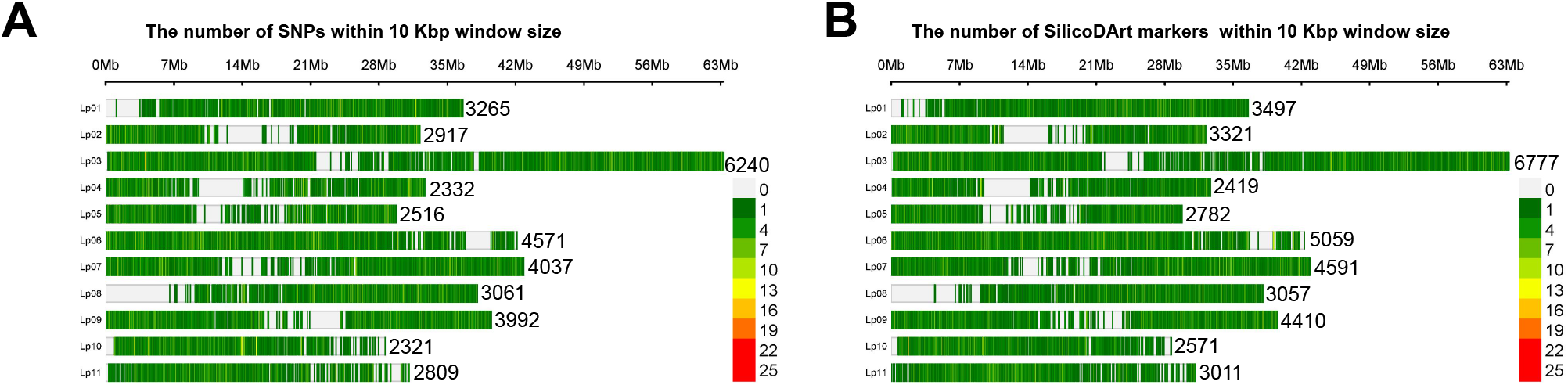
GBS polymorphism in global lablab collection: Genome-wide distribution of SNPs (**a**) and SilicoDArT (**b**) markers across the eleven chromosomes of the lablab reference genome. The total number of SNPs or SilicoDArT markers are presented beside each chromosome. Plots produced with SRplot.

**Figure S4:**
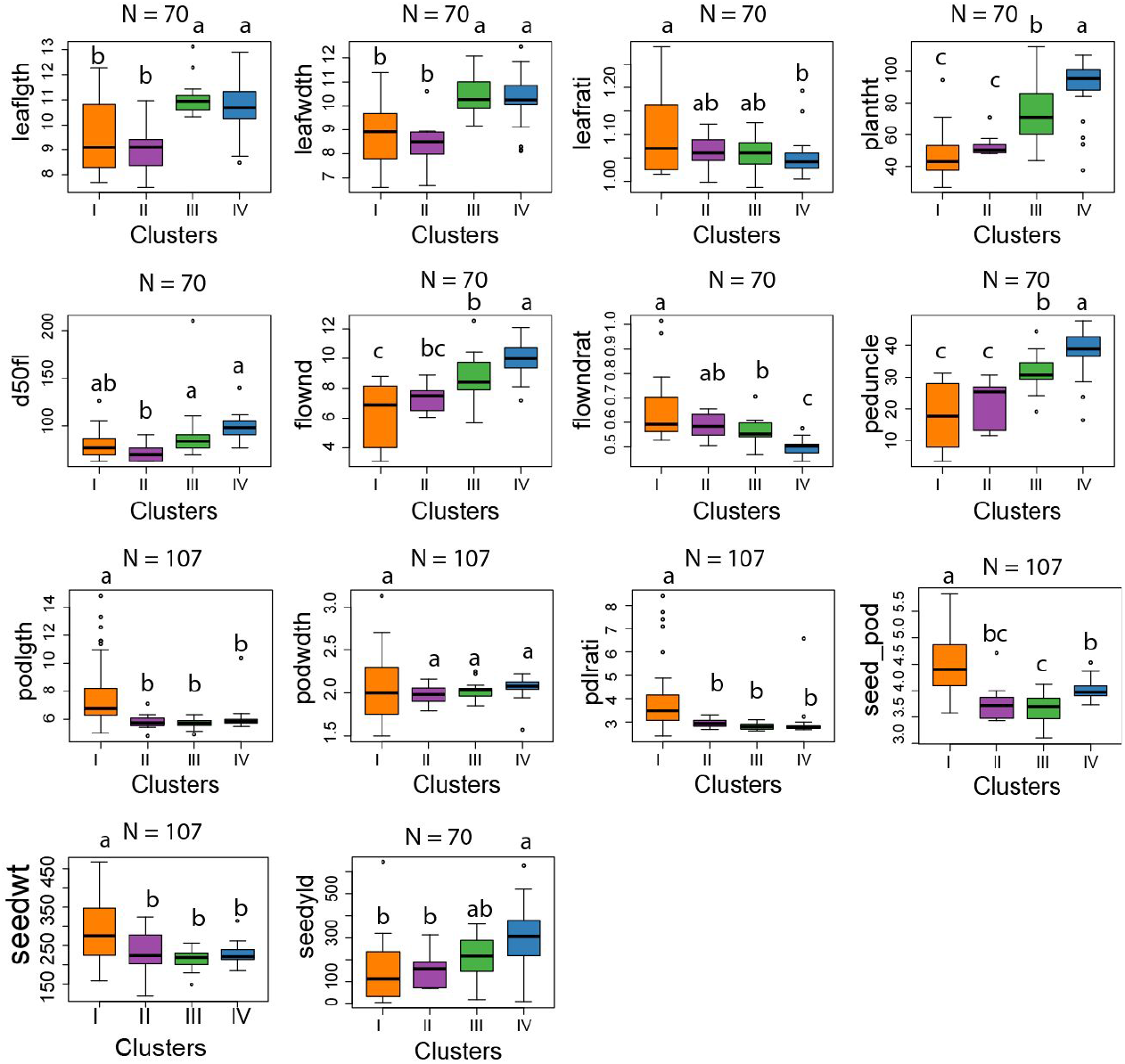
Quantitative phenotypic variation in global lablab collection. Boxplots showing phenotypic variation of different morpho-agronomic quantitative traits among the four genetic clusters identified in lablab. The colours are according to the STRUCTURE analysis with k = 4, and trait abbreviations are explained in Table S13.

**Figure S5:**
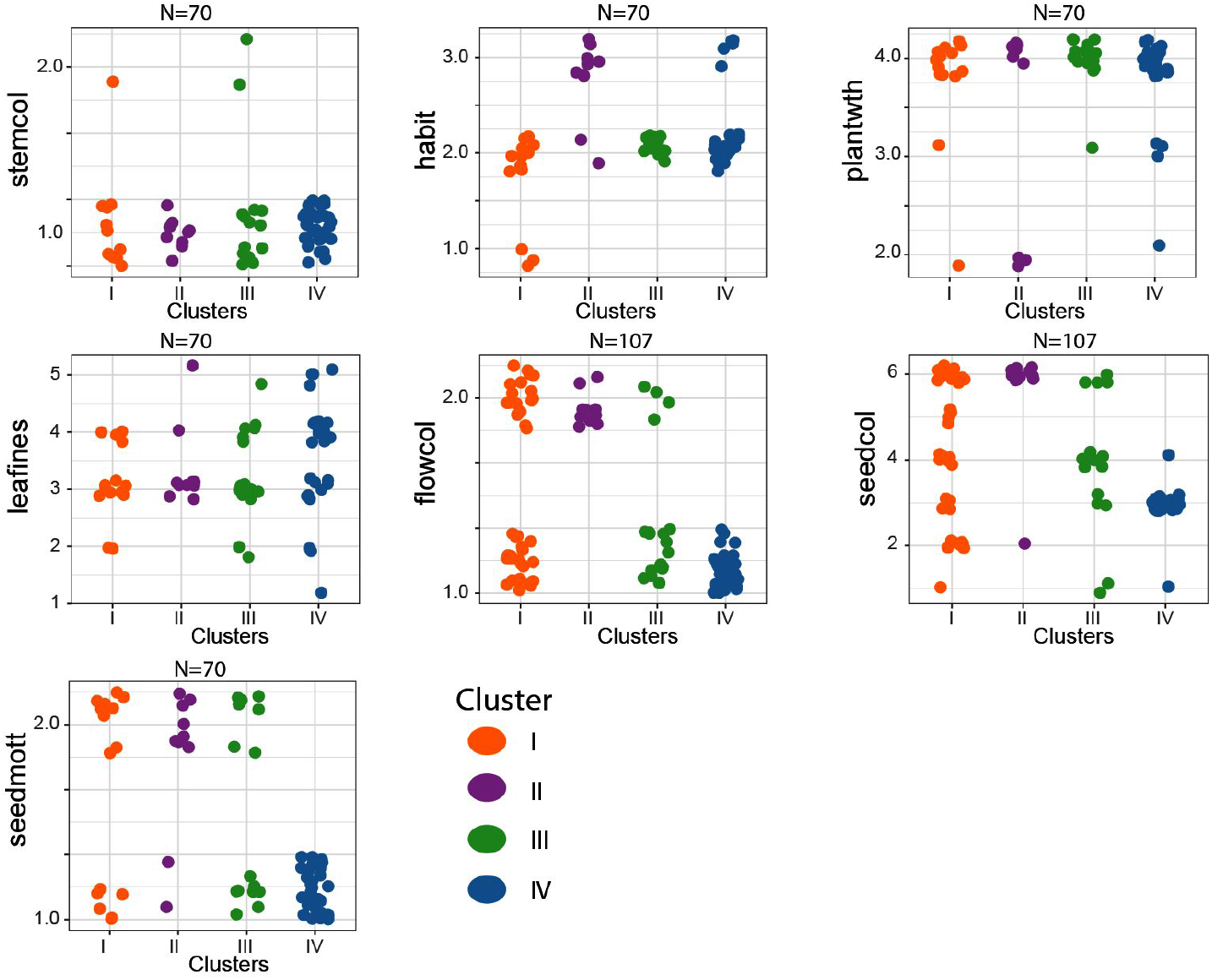
Qualitative phenotypic variation in global lablab collection. Plots showing phenotypic variation of seven qualitative traits among the four genetic clusters identified in lablab. The colours are according to the STRUCTURE analysis with k = 4, and trait abbreviations are explained in Table S14. Points are scattered if identical values are present.

**Figure S6:**
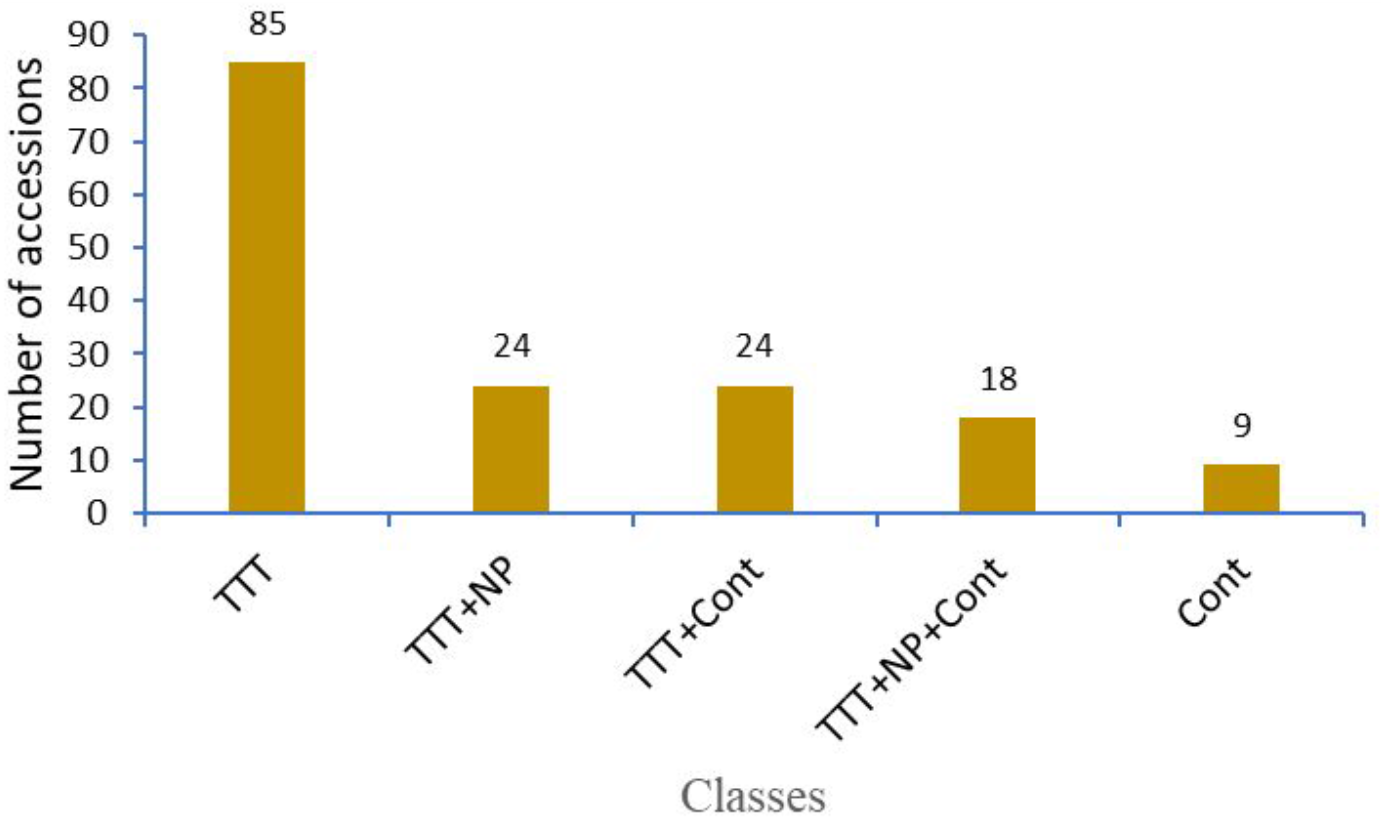
Identity-By-Descent classification of global lablab collection. Number of accessions classified as true-to-type (TTT), true-to-type and progenies (TTT+NP), true-to-type and contaminants (TTT+Cont), true-to-type and progenies and contaminants (TTT+NP+Cont), and accessions with 100% contaminants (Cont), based on a pairwise Identity-By-Descent (IBD) analysis.

## Supplementary Tables

Table S1: Summary of Nanopore Reads Statistics

Table S2: Table S2: Comparison of assembly statistics for the lablab genome based on short reads and long reads.

Table S3: Summary statistics of genes in the lablab genome.

Table S4: The number of TEs, TE families and the proportion of occupied assembly length by different classes of repeats identified and annotated in the lablab genome.

Table S5: Types, amount and proportion of tandem repeats in the lablab genome

Table S6: GO annotation of lablab-specific gene clusters

Table S7: GO annotation of gene families expanded in lablab

Table S8: Details and sequencing statistics of resequencing samples

Table S9: Population group membership

Table S10. Pairwise Fixation index (Fst) among the four major clusters (C) detected by the STRUCTURE analysis

Table S11: AMOVA showing the genetic variance among and within clusters

Table S12: Minimum, maximum and average genetic divergence (Nei’s D) between accessions within the four clusters identified by STRUCTURE.

Table S13: Results of the analysis of variance for 13 quantitative traits among the four genetic clusters.

Table S14: Results of the χ2 analysis for seven quantitative traits among the four genetic clusters. Table S15: Data on inclusive crop genomics

Table S16: Membership probability of accessions from the STRUCTURE analysis

